# Grammar acquisition in preschool children is related to white matter maturation of the dorsal language network

**DOI:** 10.1101/2025.02.23.639734

**Authors:** Cheslie C. Klein, Philipp Berger, Charlotte Grosse Wiesmann, Angela D. Friederici

## Abstract

In preschool years, children take important steps in grammar acquisition, which are essential to learning their native language. A central aspect is the acquisition of the morpho-syntactic rule system, which forms an intersection between words and sentences. In adults, rule-based linguistic processes are supported by the dorsal fiber pathway to BA44, the arcuate fascicle. This pathway matures relatively late in development, raising the question of whether it already supports grammar processes in the early preschool years, or whether early grammar acquisition is supported by different, earlier-maturing fiber pathways. In two independent samples of 3- to 5-year-old children (N = 90 and N = 30), we examined the association between the maturation of fiber pathways of the language network and children’s noun plural assignment as an index of their morpho-syntactic abilities. This revealed consistent differences between 3-year-olds and 4- to 5-year-olds. The 4- and 5-year-olds, but not 3-year-olds, showed a relation of morpho-syntax with both the dorsal pathway to BA44, supporting syntactic processes, and the dorsal pathway to BA6, supporting phonological processes in adults. Our results suggest that, in contrast to adults, preschool-aged children rely on both dorsal fiber pathways for morpho-syntax. This difference might point to different processing strategies reflecting the transition from phonology-based statistical learning to rule-based learning in grammar acquisition.

## Introduction

Grammar is at the core of human language. When acquiring their native language, young children are faced with a difficult task. Not only do they need to learn a large number of new words from their input. They also need to extract rules governing the combination of words in a sentence as well as the combination of morphological elements of a word. It is an open question how the developing brain supports these processes during language acquisition.

While words contain the meaning, grammar rules set the relation between these words in a sentence, thereby allowing the formation of complex thoughts beyond the word-level. At the word-level grammatical rules guide the combinatorial processes of morphological elements in a word. Morpho-syntax builds a bridge between the structure of the sentence and its words by modulating the grammatical features of a given word in relation to the sentence. For example, the plural form of a noun takes on a more complex structure (e.g., dog-s) than the singular noun stem (e.g., dog) by combining it with a plural morpheme (e.g., -s). Even though the combinatorial process is on the word-level, it enables the production and comprehension at the sentence-level (Harley, 2015).

During development, the ability to process rule-based regularities in auditory sequences is present in the first months of life, reflecting statistical learning of phonologically encoded regularities (Friederici et al., 2011; Marcus et al., 1999). Sentence-level syntactic rules are detected by the age of 2.5 years (Oberecker et al., 2005; Oberecker & Friederici, 2006). By 3 years of age, children can already produce and understand a variety of sentences (Kauschke, 2012; Schipke et al., 2012; Skeide et al., 2016; Strotseva-Feinschmidt et al., 2019), indicating that basic combinatorial processes seem to be in place by that age. However, milestones of complex grammar acquisition are only reached later, between the ages of 4 to 5 years and beyond (Crain & Thornton, 2012; Skeide & Friederici, 2016). Thus, the preschool years are considered as a take-off phase towards more complex grammar (Crain & Thornton, 2012; Fox & Grodzinsky, 1998; Guasti, 2002; Kauschke, 2012; Tomasello & Brooks, 1999).

At the neural level, the language network differs considerably between the developing and the adult brain. In adults, the neural language network consists of dedicated language regions in the inferior frontal cortex and the posterior temporal cortex known as Broca’s and Wernicke’s areas and the dorsally and ventrally located white matter fiber pathways interconnecting these regions (Friederici & Gierhan, 2013). In adults, grammatical processes involve BA44 in the posterior part of Broca’s area in the left inferior frontal gyrus in interaction with the posterior temporal lobe on the level of words (Bulut, 2022; Leminen et al., 2019; Meykadeh et al., 2023; Ullman, 2001) as well as sentences (Goucha & Friederici, 2015; Grodzinsky et al., 2021; Zaccarella et al., 2017). The dorsal fiber pathway that connects these two brain regions, referred to as arcuate fascicle, has been suggested to support complex grammar in adults (Friederici & Gierhan, 2013; Griffiths et al., 2013; Wilson et al., 2011) including morpho-syntactic processes (Rolheiser et al., 2011). The ventral pathway, via the inferior fronto-occipital fascicle (IFOF), is proposed to be involved in semantic processes (Fridriksson et al., 2016; Friederici & Gierhan, 2013; Rolheiser et al., 2011; Saur et al., 2008). The arcuate fascicle, dorsally connecting to BA44, can be distinguished from a second dorsal fiber bundle that connects the auditory cortex in the superior temporal lobe and BA6 in the premotor cortex. This latter pathway has been suggested to be involved in auditory-to-motor mapping as, for example, in speech repetition in adults (Fernández-Miranda et al., 2015; Friederici & Gierhan, 2013; Saur et al., 2008) and babbling in infants (Hickok & Poeppel, 2004).

Developmental brain studies have reported functional changes in the allocation of the language-related brain regions (Klein et al., 2023; Skeide et al., 2014) as well as maturational changes of the different fiber tracts (Skeide et al., 2016). A functional shift from involvement of the temporal cortex to the inferior frontal cortex has been observed between 3 and 4 years of age for sentence-level grammar processes (Klein et al., 2023). This shift towards more mature brain regions might not only underly the behavioral milestones in grammar between 3 and 4 years but might also point to alternative processing strategies for basic combinatorial processes before the age of 4 years. With respect to the fiber tracts, the dorsal pathway to BA6 and the IFOF are already detectable at birth, even though both fiber pathways still mature during childhood (Brauer et al., 2013; Perani et al., 2011). In contrast, the dorsal pathway to BA44 (i.e., the arcuate fascicle) could not be reconstructed in newborns, suggesting a structurally less mature fiber pathway compared to the other two (Perani et al., 2011). This has led to the hypothesis that early language development is predominantly supported by the dorsal pathway to BA6 and the IFOF, whereas the acquisition of complex sentence-level grammar from 4 years of age might require a more matured dorsal fiber connection to BA44 (Eichner et al., 2024; Skeide et al., 2016; Skeide & Friederici, 2016). This leaves open the question of how the maturation of the core language fiber pathways supports grammar acquisition on the word-level (i.e., morpho-syntax) in preschool-aged children.

The present study aims to clarify how the use and acquisition of morpho-syntactic rules relates to the maturation of core fiber connections of the language network in preschool years. We therefore investigated the association between developmental change in the dorsal and ventral fiber pathways of the language network and children’s grammar proficiency at the word-level during the critical preschool period of 3 to 5 years. For this, we analyzed preschooler’s morpho-syntactic performance in relation to structural properties of the two dorsal pathways targeting BA44 and BA6, and the ventral pathway (i.e., the IFOF), as well as a control tract, the corticospinal tract. This was investigated in two independent samples of 3- and 5-year-old children (Sample 1), and 3- and 4-year-old children (Sample 2), covering the ages where the shift in cortical brain regions and behavioral milestones in morpho-syntax have been observed. We hypothesized that children’s grammar abilities would be associated with indices of white matter maturation in the dorsal fiber pathway to BA44. In addition, we expected different patterns in this association between the 3-year-old versus the 4- and 5-year-old children, with younger children still relying on the well-matured dorsal pathway to BA6 or the IFOF.

## Methods

The analysis pipeline and exclusion criteria were preregistered at AsPredicted (#90877).

### Participants

For this study, MRI and behavioral data from two independent samples of monolingual German-speaking preschool children were analyzed. Sample 1 (N = 90) included 3- and 5-year-old children (3-y.o.s: N = 34, mean age = 3.63, SD = 0.24, range = 3.10 to 3.99, 12 female; 5-y.o.s: N = 56, mean age = 5.48, SD = 0.47, range = 4.01 to 6.16, 27 female). Sample 2 (N = 30) included 3- and 4-year-old children (3-y.o.s: N = 13, mean age = 3.26, SD = 0.18, range = 3.07 to 3.59, 8 female; 4-y.o.s: N = 17, mean age = 4.32, SD = 0.18, range = 4.02 to 4.58, 8 female). Children had no reported history of medical, neurological, or psychiatric disorder and no hearing or vision deficit.

These samples were subsamples from larger behavioral samples of N = 221 for the 3- and 5-year-olds of Sample 1, and N = 60 for the 3- and 4-year-olds of Sample 2. The behavioral samples included children who provided data from a standardized test battery of general language development (SETK 3-5; *Sprachentwicklungstest für drei-bis fünfjährige Kinder: SETK3–5*; Grimm, 2001), independent of whether they also underwent MRI scans. Of these, N = 88 (Sample 1) and N = 9 (Sample 2) did not participate in or aborted the MRI. Further, children were excluded if they did not provide behavioral data of any relevant SETK 3-5 subtest (Sample 1: N = 1) or showed an indication for a speech development delay in the SETK 3-5 (T-value < 35; Sample 1: N = 8, Sample 2: N = 2). Children were excluded from the MRI analyses if they received a later diagnosis of ADHD or developmental dyslexia (Sample 1: N = 16) or did not provide handedness data (Sample 1: N = 4, Sample 2: N = 10). After MRI preprocessing, children were further excluded if they showed severe motion artifacts in the MRI scans (see *MRI preprocessing*; Sample 1: N = 10, Sample 2: N = 6), or for which brain extraction, estimation of gradient direction or tractography failed (see *MRI preprocessing*; Sample 1: N = 5, Sample 2: N = 3). For the standardization of the behavioral scores, children were further excluded from the initial behavioral sample if they did not provide data of all SETK 3-5 subtests (Sample 1: N = 18). This resulted in N = 194 3- and 5-year-old children for standardization within Sample 1 and N = 58 3- and 4-year-olds within Sample 2.

Parental consent was obtained for all children and the study was approved by the Ethics Committee at the Faculty of Medicine of the University of Leipzig (number of approval; Sample 1: 320-11-26092011, Sample 2: 090/12-ff). Data of both samples have previously been analyzed separately with regard to other research questions (Sample 1: Eichner et al., 2024; Kuhl et al., 2020, 2021; Liebig et al., 2020; Qi et al., 2021; Sample 2: Berger et al., 2021; Grosse Wiesmann et al., 2020; Grosse Wiesmann, Friederici, et al., 2017; Grosse Wiesmann, Schreiber, et al., 2017; Klein et al., 2023).

### Behavioral data

#### Morpho-syntactic ability

The SETK 3-5 is a comprehensive diagnostic tool designed to test 3- to 5-year-old children’s general language development in the domains of language comprehension, production, and memory (Grimm, 2001). To test children’s grammar abilities on the word-level, we selected and preregistered the morpho-syntactic word production task (SETK 3-5 subtest ‘Morphologische Regelbildung’; Grimm, 2001), which determines the acquisition level of the morpho-syntactic rule system for German plural formation.

In this task children are asked to name the plural of given nouns. For this, children are first shown a picture of an object or animal that is described using the singular form of a noun (e.g., “Schau mal, hier ist ein Auto.” [engl. “Look, here is a car.”]; Grimm, 2001). Then, a picture with three of the same objects or animals is shown and the child is asked to produce the plural form of the given word (e.g., “Hier kommen noch mehr dazu. Hier sind drei…?” [engl. “Here are some more. Here are three…?”]; Grimm, 2001). Children are familiarized with the task with one practice item and then tested on ten common real nouns, which follow the various dominant plural rules of the German noun system (for further description of the German plural system see *Supplementary Methods*). The 4- to 5-year-old children were presented, in addition, with eight pseudo words similar to the Wug Test from Berko (1958) (for a list of the items see *Supplementary Table 1*; Grimm, 2001).

Children’s responses for each item were rated depending on whether a plural rule was used, and which one. A score of 2 was given for the correct plural form, of 1 for the use of a different but incorrect plural rule (e.g., “Fisch-en” instead of “Fisch-e”; Grimm, 2001). A score of zero was given for an incorrect response, for example, repetition of the singular form or a plural noun with significant changes of the word stem (e.g., “Gratel-en” instead of “Gabel-n”; Grimm, 2001), or no response at all. In the present study, we used raw values of the morpho-syntactic word production task rather than normalized scores provided by the SETK 3-5, as we were interested in relating individual developmental differences to brain structure. To account for the differences in item structure among 3- and 4- to 5-year-olds, we z-transformed these raw scores within each age group. The standardization was carried out independently within the full behavioral datasets of each sample (Sample 1: N = 194, Sample 2: N = 58). The resulting morpho-syntax score was used for the brain-behavior analyses.

In addition, we preregistered a syntactic comprehension and production score from two further subtests of the SETK 3-5 to test for children’s grammar ability on the sentence-level, which we report in the Supplementary Material.

#### Psychometrics

Children’s non-verbal IQ was assessed using the Wechsler Preschool and Primary Scale of Intelligence (WPPSI-III; Wechsler, 2009) in Sample 1, and the Kaufman Assessment Battery for Children (Kaufman ABC; Melchers & Preuß, 2003) in Sample 2. Children’s non-verbal IQ scores were within the typical range for the respective age: For Sample 1, the standard scores of the non-verbal scale were used (mean = 98.33, SD = 13.30), and for Sample 2, the mean scale scores were used (mean = 10.26, SD = 1.34). In both samples, children’s degree of handedness was evaluated with the German version of the Edinburgh Handedness Inventory (Oldfield, 1971).

### MRI data acquisition

MRI data were obtained on a 3T Siemens scanner (Siemens MRT Trio series) equipped with a 12-channel head coil for Sample 1 and a 32-channel head coil for Sample 2.

In Sample 1, diffusion-weighted MRI (dMRI) data were acquired using the optimized monopolar Stejskal-Tanner EPI sequence (Morelli et al., 2010) along the anterior-to-posterior phase encoding (PE) direction (voxel size = 1.86*×*1.86*×*1.9 mm, TR = 8,000 ms, TE = 83.0 ms, b-value = 1,000 s/mm^2^, 60 directions, GRAPPA 2). A field map was acquired after completion of each 10 images. In addition, one diffusion-weighted image and a field map was acquired along the posterior-to-anterior PE direction. In Sample 2, dMRI data were acquired using the multiplexed echo planar imaging (EPI) sequence (Feinberg et al., 2010) with a spatial resolution of 1.9 mm isotropic (TR = 4,000 ms, TE = 75.4 ms, b-value = 1,000 s/mm^2^, 60 directions, GRAPPA 2). After the dMRI scan, a field map was acquired as an anatomical reference.

High-resolution 3D T1-weighted MRI data were further acquired in both samples using the MP2RAGE sequence (Marques et al., 2010) (Sample 1: voxel size = 1.3 mm isotropic, TR = 5,000 ms, TE = 2.82 ms, TI_1_ = 700 ms, α_1_ = 4°, TI_2_ = 2,500 ms, α_2_ = 5°; Sample 2: voxel size = 1.2*×*1.0*×*1.0 mm, TR = 5,000 ms, TE = 3.24 ms, TI_1_ = 700 ms, α_1_ = 4°, TI_2_ = 2,500 ms, α_2_ = 5°).

A summary table of the MRI parameters applied in the two samples can be found in the Supplementary Material (see *Supplementary Table 2*).

### MRI data analysis

MRI preprocessing and analyses were performed independently within each of the two samples, unless otherwise stated.

#### MRI preprocessing

MRI preprocessing of the dMRI data included the following steps (Eichner et al., 2024; Fan et al., 2022). Estimation of the susceptibility-induced off-resonance field was done using FSL’s *topup* tool to correct for distortions (Andersson et al., 2003; Smith et al., 2004). The underlying noise distribution was then used to correct for non-Gaussian noise biases in the data (St-Jean et al., 2020). Further, data was denoised and Gibbs ringing artefacts were removed using MRtrix3’s *dwidenoise* and *mrdegibbs* (Kellner et al., 2016; Tournier et al., 2019; Veraart et al., 2016). Motion correction was applied with FSL’s *eddy* (Andersson & Sotiropoulos, 2016). We additionally performed signal drift correction using a second-order polynomial fit to the data without diffusion-weighting (Vos et al., 2017).

T1-weighted MRI data were preprocessed with the following steps. Background noise was removed by combining the two inversion time images (O’Brien et al., 2014). The fsl_anat pipeline was then used to reorient and register the data to the MNI space, perform brain extraction, and segment the brain tissues (http://fsl.fmrib.ox.ac.uk/fsl/fslwiki/fsl_anat). Then, the data was aligned to ACPC using the Automatic Registration Toolbox (https://www.nitrc.org/projects/art). Finally, diffusion data were aligned to the processed T1-weighted images following the *dtiInit* pipeline from the VISTASOFT package (https://github.com/vistalab/vistasoft).

After preprocessing of the diffusion- and T1-weighted MRI data, we performed several quality checks on the processed data. First, we used FSL’s *eddy QC* tool to perform quality assessment of diffusion data on the single-subject level (Bastiani et al., 2019). Based on the output of this tool, we excluded children with large head movement values, more than 2 SDs from each independent sample’s mean movement. Second, a diffusion tensor model was fitted at each voxel using FSL’s *dtifit* and the eigenvectors of prominent tracts as the superior longitudinal fascicle, corpus callosum, and corticospinal tract were visually checked on the FA images. Third, successful brain extraction was visually checked, both for the diffusion- and T1-weighted MRI data.

#### Automated tractography pipeline

To reconstruct the left dorsal and ventral language-related fiber pathways and the corticospinal tract as a control from the preprocessed dMRI data, we used the python-based version of the Automated Fiber Quantification (pyAFQ 0.12) software (Kruper et al., 2021; Yeatman et al., 2012). This pipeline utilizes several steps to generate well-defined and anatomically plausible fiber pathways. Further, it results in global measures such as the streamline count of each pathway, as well as so-called tract profiles of local assessments of microstructural diffusion properties along each pathway (Kruper et al., 2021; Yeatman et al., 2012).

We performed the following steps as provided by the pyAFQ pipeline. The distribution of fiber orientation at each voxel was estimated using the Constrained Spherical Deconvolution (CSD) model (Tournier et al., 2004). A whole-brain probabilistic tractography was then performed with maximum turning angle = 30°, minimum streamline length = 10Lmm, maximum length = 1000 mm, and step-size = 0.5 mm.

The resulting tractograms were then segmented into the three language-related fiber pathways of interest and the additional control tract by using predefined endpoint and waypoint ROIs, based on prior anatomical knowledge. For the endpoint ROIs, regions from the AAL atlas (Tzourio-Mazoyer et al., 2002) were selected to define the cortical termination points of each fiber pathway. Waypoint ROIs were defined in MNI space and then registered to children’s native space using a non-linear transformation to further guide the segmentation to result in anatomically plausible pathways (Wakana et al., 2007). Within the pyAFQ pipeline, endpoint and waypoint ROIs for over 20 major fiber pathways are provided, including the arcuate fascicle, IFOF, and corticospinal tract. To disentangle the dorsal pathway to BA44 from the one to BA6, we modified the default segmentation pipeline for the arcuate fascicle (see *Supplementary Figure 1*). For this, we used the default endpoint ROI in the temporal lobe but defined BA44 and BA6 as endpoint ROIs in the frontal cortex, respectively. Further, we used the default waypoint ROIs of the arcuate fascicle for both dorsal fiber pathways. For the dorsal pathway to BA6, we additionally defined an exclusion ROI in BA44 to disentangle streamlines of both dorsal fiber pathways, and a second exclusion ROI to ensure that no fibers would branch into the corticospinal tract. To segment the ventral fiber pathway, corresponding to the IFOF and the corticospinal tract as control tract, we relied on the default endpoint and waypoint ROIs provided by pyAFQ.

The segmented fiber pathways were then further refined by comparing each candidate fiber to a probability map (Hua et al., 2008) and discarding fibers passing low probability regions. By default, pyAFQ provides probability maps for the arcuate fascicle, which was used for both dorsal fiber pathways, for the IFOF, and corticospinal tract. Outlier fibers that deviated substantially from the core of the respective fiber pathway were additionally removed to result in a compact fiber bundle. To identify the fiber core, each pathway was clipped at the endpoint ROIs, resampled into 100 equidistant nodes along the pathway and the mean location of each node was calculated. The resulting bundles of each child were visually inspected to ensure biologically plausible identification of each fiber pathway. Fiber bundles of the two dorsal fiber pathways were excluded if streamlines followed anatomically implausible turns, or if the frontal endpoints of the bundles did not terminate in distinct areas of BA44 or BA6 (for examples see *Supplementary Figure 2*). The visual inspection led to the exclusion of N = 4 of the dorsal pathway to BA44, N = 2 of the dorsal pathway to BA6, and N = 1 of the corticospinal tract in Sample 1, and N = 1 of the IFOF Sample 2.

Diffusion properties (i.e., fractional anisotropy [FA], mean diffusivity [MD], and radial diffusivity [RD]) were quantified at each node along each fiber pathway using a weighted approach as described by Yeatman et al. (2012). This resulted in a so-called tract profile that was used for further analysis. In addition, total streamline count of each fiber pathway was calculated.

We additionally reconstructed the homologous dorsal and ventral language pathways and the corticospinal tract as a control in children’s right hemisphere. These exploratory analyses are reported in the Supplementary Material.

#### Statistical analysis

To estimate the relation between children’s morpho-syntactic ability and structural properties of the language-related fiber pathways and the control tract, we fit multiple linear regressions using the lm() function from the stats package included in R (R Core Team, 2023) using RStudio (RStudio Team, 2020). As preregistered, we used the morpho-syntax scores as the predictor and the brain structural measures (i.e., total streamline count, FA, MD, and RD) as dependent variables in each model. We included both age groups in the analyses, with age group as factor and tested for a main effect and an interaction between age group and children’s morpho-syntax scores. These analyses were performed independently in the two samples. Significant interactions with age group were followed up in each age group separately. In addition, for the purpose of comparison, we also explored language scores with a nonsignificant age interaction in the separate age groups. In all models, we controlled for sex, non-verbal IQ, handedness, and estimated intracranial volume (eTIV) to examine the specificity of the effects. As Sample 1 of the 3- and 5-year-olds consisted of children with and without a family history of dyslexia, we included family history as an additional covariate in the analyses of this dataset. In all models testing for a main effect, we performed one-sided tests as streamline count and FA increase with maturation and higher cognitive function, while MD and RD decrease (Lebel & Deoni, 2018). Since streamline count constitutes one value per fiber pathway, the alpha-level was set at *p* = 0.05 for the models including streamline count. In the node-based approach (i.e., for FA, MD, and RD), multiple linear regressions were performed pointwise along 100 segments (i.e., nodes) of each fiber pathway. We therefore applied cluster-wise correction for multiple comparisons at *p* = 0.05 using non-parametric permutation-based correction (Nichols & Holmes, 2002) with a further Bonferroni correction for the N = 3 microstructural indices. Statistics for the cluster-wide relationship between these indices and morpho-syntax scores were calculated on the mean FA, MD, or RD of the respective significant cluster. We winsorized all language data and brain maturational indices limiting extreme values to the 95th percentile prior to conducting analyses.

To relate brain maturational indices of the language-related fiber pathways to preschooler’s grammar performance on the sentence-level, we performed similar analyses on children’s syntactic comprehension and production scores as reported in the Supplementary Material.

## Results

To investigate the relation between preschooler’s abilities in morpho-syntactic processing and brain structural maturation of language-related fiber pathways, we reconstructed two dorsal fiber pathways, one targeting BA44 and one targeting BA6, as well as the ventral pathway (i.e., IFOF), and the corticospinal tract as a control tract (see Fig. 1). In a preregistered procedure, we then related their brain maturational indices (i.e., total streamline count and local measures of FA, MD, and RD along 100 nodes of each pathway) to children’s morpho-syntactic abilities, which were assessed using a noun plural assignment task. We examined this relation in two independent samples of 3- and 5-year-old (Sample 1), and 3- and 4-year-old (Sample 2) children. All effects were independent of sex, non-verbal IQ, handedness, eTIV, and, in Sample 1, additionally independent of family history of dyslexia. For models with streamline count, the significance threshold was set at *p* = 0.05. For node-based models, multiple comparison correction was applied at *p* = 0.05 with a further Bonferroni-correction to correct for N = 3 brain maturational indices.

**Figure 1:**
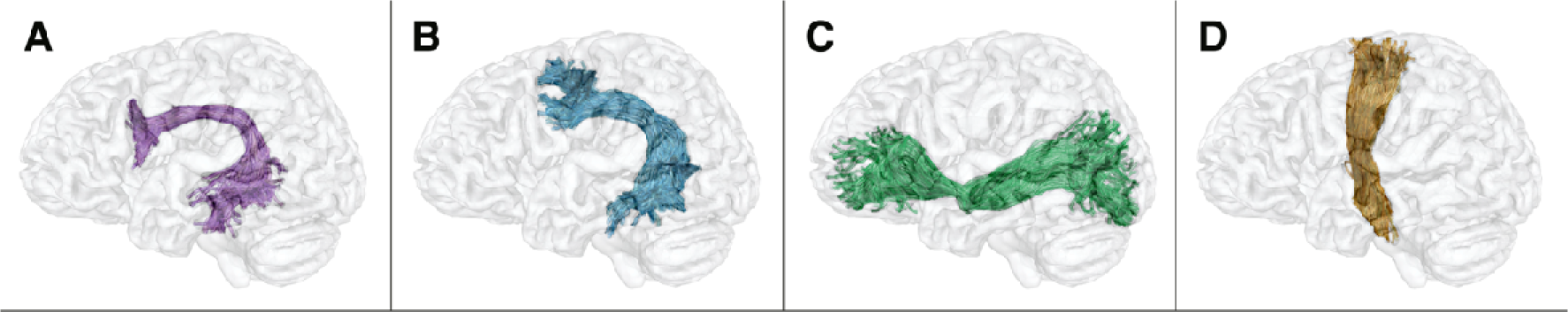
Example of the segmented and refined language fiber pathways – a) the dorsal pathway to BA44 (purple), b) the dorsal pathway to BA6 (blue), the c) ventral pathway (green) – and d) the corticospinal tract as control tract (orange).

In both samples, younger children at the age of 3 years differed from older children aged 4 and 5 years. In Sample 1, there was no significant main effect of children’s morpho-syntax scores on their brain maturational measures in the reconstructed pathways, but a significant interaction with age group. This age interaction was found in the dorsal pathway to BA6 (anterior part: RD, node size = 15, node range = 17-31, β = −0.03, SE = 0.01, *t*(78) = −3.076, *p* = 0.003) and in the ventral pathway (streamline count, β = 413.4, SE = 181.5, *t*(80) = 2.278, *p* = 0.025; central part: FA, node size = 14, node range = 39-52, β = 0.032, SE = 0.001, *t*(80) = 4.400, *p* < 0.001; RD, node size = 12, node range = 40-51, β = −0.031, SE = 0.01, *t*(80) = - 3.166, *p* = 0.002), but not the dorsal pathway to BA44. No main effect or interaction with age was found in the corticospinal tract serving as control tract. To follow-up on these interactions, we analyzed the two age groups, i.e. the 3- and 5-year-olds, separately. This revealed that the interactions were driven by the 5-year-old children who showed a significant relation between their morpho-syntax scores and brain maturational measures in the dorsal pathway to BA6 (anterior part: FA, node size = 14, node range = 21-34, β = 0.0230, SE = 0.008, *t*(47) = 3.069, *p* = 0.002; RD, node size = 16, node range = 18-33, β = −0.0197, SE = 0.007, *t*(47) = −2.832, *p* = 0.003; see Fig. 2a) and the ventral pathway (streamline count, β = 322.0, SE = 136.3, *t*(48) = 2.363, *p* = 0.011; central part: FA, node size = 17, node range = 42-58, β = 0.021, SE = 0.065, *t*(48) = 3.241, *p* = 0.001). No effect was found in the dorsal pathway to BA44 or the control tract in the 5-year-olds. The 3-year-olds did not show any significant relation in either of the three language-related fiber pathways or the control tract.

**Figure 2:**
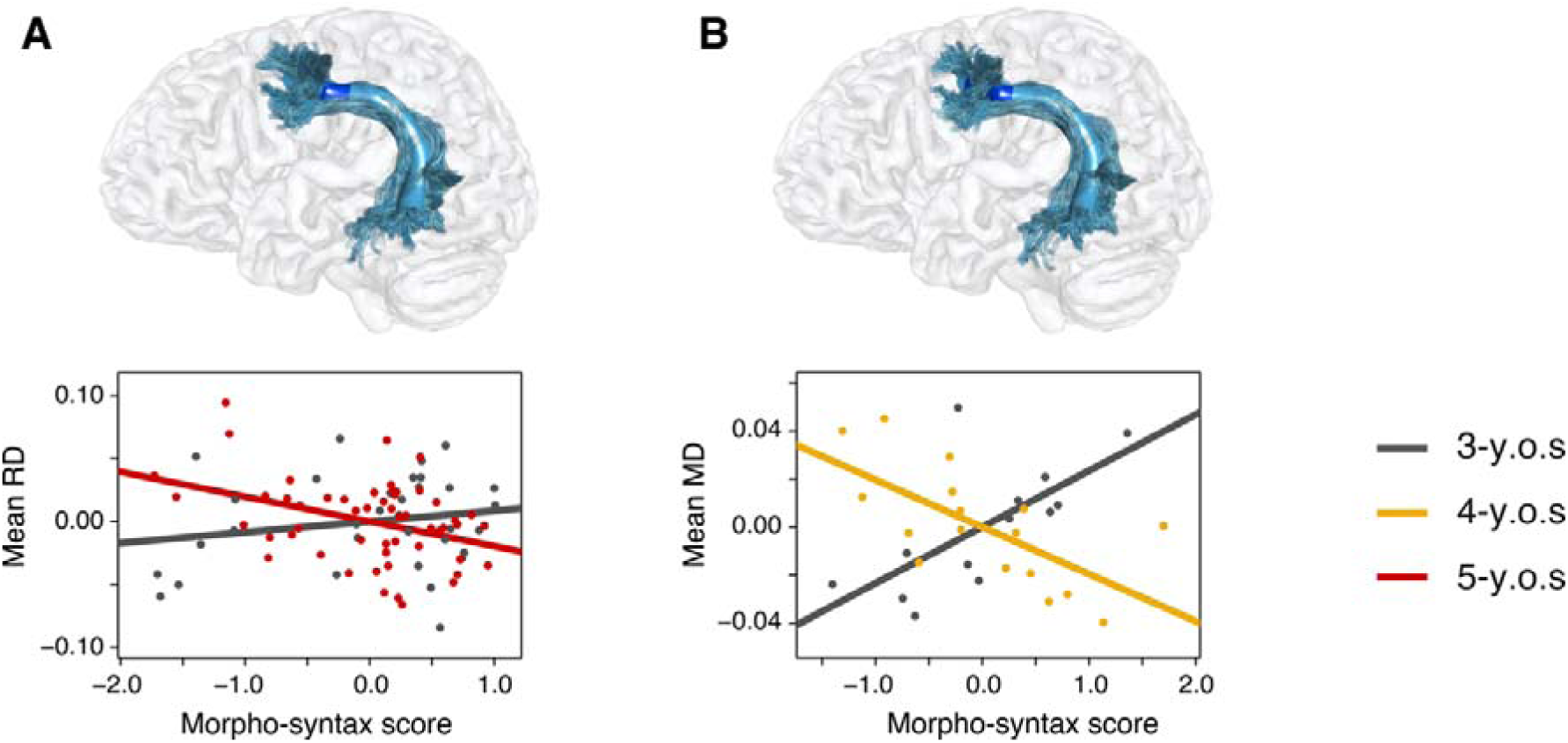
BA6 targeting dorsal pathway. Linear partial correlation of preschooler’s morpho-syntactic abilities and brain structural measures within the significant clusters of the dorsal pathway to BA6 (blue) in (A) Sample 1 and (B) Sample 2. Cluster-wise correction was applied at *p* = 0.05 with a further Bonferroni-correction of N = 3. Residuals of the partial correlation of the morpho-syntax score and the mean values of the brain maturational measures within the significant clusters are plotted. (A) Significant relation of 5-year-old’s radial diffusivity (RD) and morpho-syntax scores in Sample 1 (3-y.o.s: *r_p_* = 0.19, *p* = 0.33, gray; 5-y.o.s: *r_p_*= −0.38, *p* = 0.007, red). (B) Significant relation of 4-year-old’s mean diffusivity (MD) and morpho-syntax scores in Sample 2 (3-y.o.s: *r_p_* = 0.67, *p* = 0.05, gray; 4-y.o.s: *r_p_* = −0.67, *p* = 0.013, yellow).

In Sample 2, again no main effect of children’s morpho-syntax scores on their language-related pathways was found. Further, we replicated the interaction with age group in the dorsal pathway to BA6 (part terminating in the temporal lobe: MD, node size = 28, node range = 73-100, β = −0.031, SE = 0.009, *t*(22) = −3.568, *p* = 0.002). Additionally, however, we found an age interaction in the dorsal pathway to BA44 (posterior part: MD, node size = 20, node range = 71-90, β = −0.031, SE = 0.01, *t*(22) = −3.202, *p* = 0.004; anterior part: RD, node size = 20, node range =18-37, β = −0.045, SE = 0.011, *t*(22) = −4.058, *p* < 0.001). No interaction with age was found in the ventral pathway, nor the control tract. Follow-up analyses separated by age group revealed that these interactions were again driven by the older age group. That is, the 4-year-old children, but not the 3-year-olds, showed a significant relation in the dorsal pathway to BA6 (anterior part: MD, node size = 20, node range = 10-29, β = −0.020, SE = 0.007, *t*(11) = −2.965, *p* = 0.006; see Fig. 2b) and in the dorsal pathway to BA44 (anterior part: MD, node size = 27, node range = 7-33, β = −0.016, SE = 0.004, *t*(11) = - 4.172, *p* < 0.001; RD, node size = 11, node range = 27-37, β = −0.023, SE = 0.007, *t*(11) = −3.240, *p* = 0.004). In line with the absence of an interaction, no significant effect was found in the ventral pathway or the corticospinal tract in either of the two age groups.

In order to test whether these correlations between behavior and white matter fiber tracts were specific for the left hemisphere, we also analyzed the corresponding white matter fiber tracts in the right hemisphere. For the right-hemispheric homologous fiber pathways, we found no main effect, but a significant age interaction driven by the older age groups in the right ventral pathway in Sample 1 and in the right dorsal pathway to BA6 in Sample 2 for children’s morpho-syntax scores (see *Supplementary Results* for more details). No interaction or main effect was found for the right dorsal pathway to BA44 in either sample.

Further, to test whether the interaction with age found in our analyses could be due to the fact that the noun plural assignment task in 3-year-olds only contained real nouns, while in 4- and 5-year-olds additionally contained pseudo nouns, an analysis only including the real nouns was conducted. This was done as the generation of plural forms of real nouns and pseudo nouns might be based on different strategies since the plural forms of pseudo nouns cannot be retrieved from the mental lexicon but requires rule application. Thus, in an exploratory analysis, we divided the items presented to the 4- and 5-year-olds into real and pseudo nouns. The summed score of real noun items were standardized within both age groups in each sample independently, i.e. the 3- and 5-year-olds of Sample 1, and the 3- and 4-year-olds of Sample 2. We then tested for a relation between children’s real noun morpho-syntax scores and maturational indices in the language pathways and the control tract.

For children’s real noun morpho-syntax scores, as before, no significant main effect but significant interactions with age group were found in both samples. In Sample 1, the interaction was found in the dorsal pathway to BA6 (anterior part: MD, node size = 20, node range = 7-26, β = −0.025, SE = 0.007, *t*(77) = −3.498, *p* < 0.001; RD, node size = 18, node range =16-33, β = −0.041, SE = 0.011, *t*(77) = −3.781, *p* < 0.001), in the dorsal pathway to BA44 (anterior to central part: FA, node size = 9, node range = 29-37, β = 0.033, SE = 0.01, *t*(75) = 3.171, *p* = 0.002; RD, node size = 17, node range =28-44, β = −0.041, SE = 0.012, *t*(75) = −3.456, *p* < 0.001), and the ventral pathway (streamline count, β = 453.4, SE = 213.0, *t*(79) = 2.128, *p* = 0.036; central part: FA, node size = 13, node range =40-52, β = 0.034, SE = 0.009, *t*(79) = 3.762, *p* < 0.001). As before, follow-up analyses revealed that the present interactions were driven by the older age group. The 5-year-olds, but not the 3-year-olds, showed an effect in the dorsal pathway to BA6 (anterior part: RD, node size = 15, node range = 18-32, β = −0.03, SE = 0.009, *t*(47) = −3.348, *p* < 0.001) and in the dorsal pathway to BA44 (anterior to central part: RD, node size = 20, node range = 25-44, β = −0.028, SE = 0.009, *t*(45) = −3.045, *p* = 0.002; see Fig. 3a). No effect was found for the ventral pathway in either of the two age groups.

**Figure 3:**
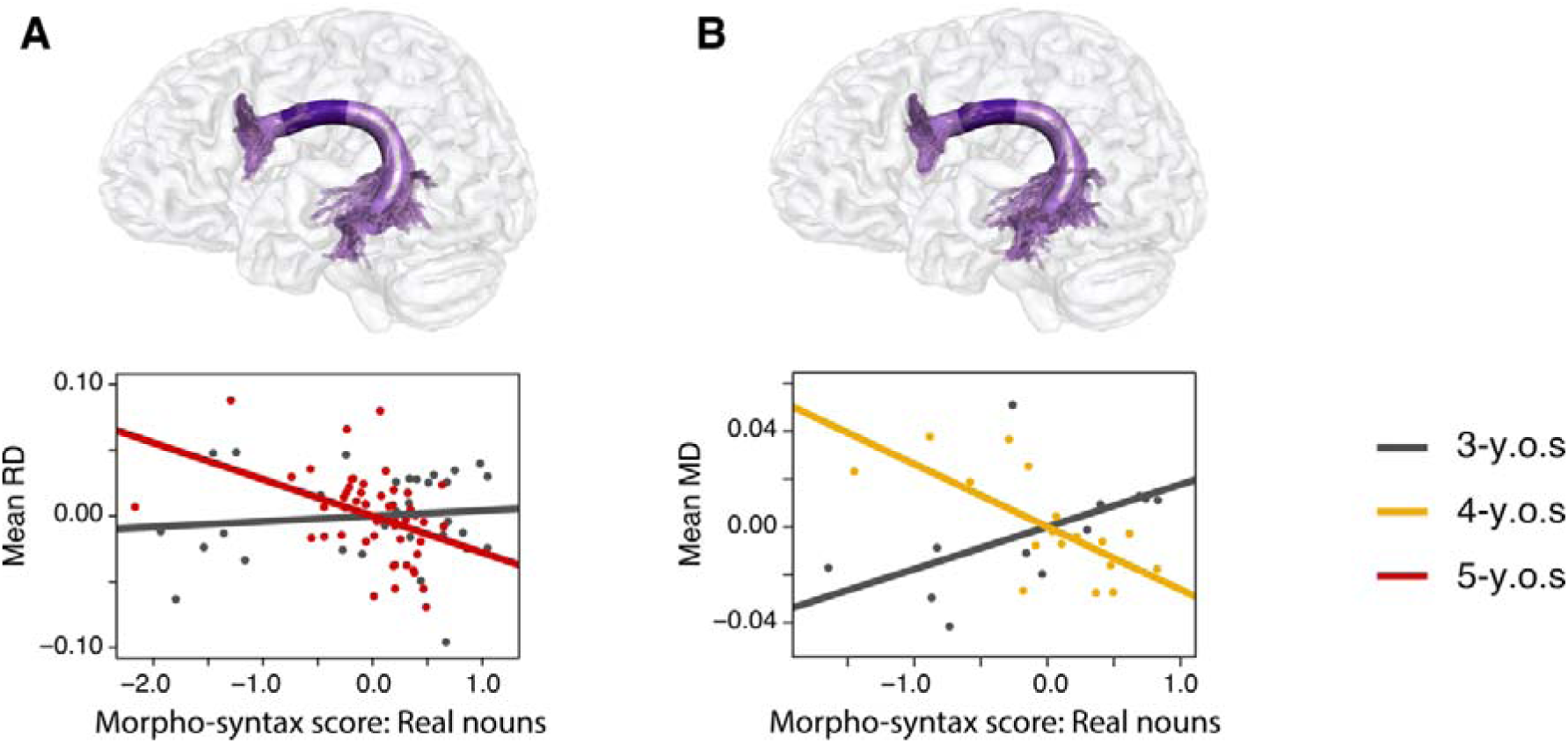
BA44 targeting dorsal pathway. Linear partial correlation of preschooler’s morpho-syntactic abilities tested with real nouns and brain structural measures within the significant clusters of the dorsal pathway to BA44 (purple) in (A) Sample 1 and (B) Sample 2. Cluster-wise correction was applied at *p* = 0.05 with a further Bonferroni-correction of N = 3. Residuals of the partial correlation of the morpho-syntax score tested with real nouns and the mean values of the brain maturational measures within the significant clusters are plotted. (A) Significant relation between 5-year-old’s real noun morpho-syntax scores and radial diffusivity (RD) in Sample 1 (3-y.o.s: *r_p_* = 0.12, *p* = 0.59, gray; 5-y.o.s: *r_p_* = −0.41, *p* = 0.004, red).(B) Significant relation between 4-year-old’s real noun morpho-syntax scores and mean diffusivity (MD) in Sample 2 (3-y.o.s: *r_p_* = 0.61, *p* = 0.08, gray; 4-y.o.s: *r_p_* = −0.70, *p* = 0.007, yellow).

In Sample 2, the interaction was also located in the dorsal pathway to BA6 (anterior to central part: FA, node size = 11, node range = 35-45, β = 0.075, SE = 0.018, *t*(22) = 4.121, *p* < 0.001; MD, node size = 34, node range = 11-44, β = −0.045, SE = 0.013, *t*(22) = −3.368, *p* = 0.003; RD, node size = 14, node range =32-45, β = −0.073, SE = 0.016, *t*(22) = −4.518, *p* < 0.001) replicating the results of Sample 1. No significant interaction was found in the dorsal pathway to BA44 or the ventral pathway in this sample. Following-up on this interaction, 4-year-old children did not show a significant effect in the dorsal pathway to BA6, but a significant relation in the dorsal pathway to BA44 (anterior part: MD, node size = 17, node range = 27-43, β = −0.026, SE = 0.008, *t*(11) = −3.296, *p* = 0.004; see Fig. 3b). No effect was found for the ventral pathway. No significant effect for the real noun morpho-syntax scores was found in the 3-year-olds in either sample. Moreover, no main effect or interaction was found for the control tract. These results indicate that the age difference between the 3-year-olds and the older children found for the entire set of nouns was not driven by different subsets of the experimental material.

Analyses for the pseudo noun plurals can be found in the Supplementary material.

## Discussion

The present paper investigated the relation between grammar acquisition at the morpho-syntactic level and brain maturation at preschool age. Children between 3 and 5 years of age are known to undergo major behavioral improvements in their morpho-syntactic performance (Kauschke, 2012). However, the dorsal fiber pathway to BA44 supporting grammar in adults matures relatively late during development (Brauer et al., 2013; Skeide & Friederici, 2016). This raises the question of whether this fiber pathway already supports morpho-syntax acquisition in the early preschool years, or whether earlier maturing brain structures take on this role. We investigated this question in two independent samples of 3- and 5-year-old children (Sample 1), and 3- and 4-year-old children (Sample 2). Our analysis showed consistent differences in the association between the maturation of language fiber pathways and children’s morpho-syntactic abilities between the 3-year-olds and the older age groups. In both samples, the older age group showed a relation of their morpho-syntax scores with the anterior part of the dorsal pathway to BA6. In an exploratory analysis, in which we analyzed this relation in real nouns only, we further found a consistent additional relation of the dorsal pathway to BA44 with morpho-syntax abilities in the 4- and 5-year-olds, but not in the 3-year-olds. These results reveal a clear difference of the involvement of the dorsal pathways BA6 and BA44 in the older compared to the younger children.

In the adult brain, the dorsal pathway to BA6 has been found to support auditory-to-motor mapping and phonological processes as in speech repetition (Saur et al., 2008). In the developing brain, this fiber pathway is already detectable in newborns, suggesting a relatively mature white matter structure at birth (Perani et al., 2011). Thus, it was hypothesized that early language acquisition might be primarily supported by the dorsal pathway to BA6 (Skeide & Friederici, 2016). Our findings that children’s morpho-syntactic abilities, which reflect rule-based processes at the word-level, were associated with this fiber pathway are in line with this hypothesis. Further, the dorsal pathway to BA6 might support the detection of statistical regularities in phonological sequences. From behavioral studies, we know that statistical learning is crucial for language acquisition in general and for grammar development in particular (Saffran, 2020). The German plural rule system relies on the phonology of a given noun, although not exclusively (Köpcke, 1988). The present association with the dorsal pathway to BA6 with children’s morpho-syntactic abilities might point to a phonology-based statistical acquisition strategy. Thus, in contrast to adults, children might strongly rely on learning statistical regularities to acquire grammatical rules, be it on the sentence-level (Friederici et al., 2011) or the word-level as in the present study, supported by the dorsal pathway to BA6.

In addition, we found a consistent relation between 4- and 5-year-olds’ real noun morpho-syntax scores and the dorsal pathway to BA44, which was also observed for the 4-year-old’s overall morpho-syntax scores. In the mature brain, this pathway supports rule-based language processes, as found both in adult and patient studies (Friederici & Gierhan, 2013; Griffiths et al., 2013; Wilson et al., 2011). In children, the maturation of the dorsal pathway to BA44 is related to sentence processing abilities from 4 years of age (Skeide et al., 2016). Preschool children evidently build up a rule system during the acquisition of plural nouns (Kauschke, 2012). This is apparent in the overgeneralization errors children produced that mainly follow frequent plural rules (see *Supplementary Figure 3*; Kauschke et al., 2011; Laaha, 2011; Szagun, 2001). Thus, maturation of the dorsal pathway to BA44 might not only support the acquisition of complex sentences (Skeide et al., 2016) but also of rule-based processes on the word-level.

To summarize, we found a consistent association between 4- and 5-year-old children’s abilities to apply morpho-syntactic rules and the maturation of both dorsal pathways (to BA44 and to BA6) across two independent samples. In adults, the dorsal connection between BA44 and the temporal cortex is crucial for rule-based processes, such as morpho-syntax (Meykadeh et al., 2023; Rolheiser et al., 2011). Against this background, our findings suggest that children may employ both statistical and rule-based learning strategies to acquire grammar in the early preschool years, supported by the maturation of the dorsal fiber pathways to BA6 and BA44.

### Limitations

While the main findings were consistent across two independent samples – the association of the dorsal pathway to BA6 with children’s morpho-syntax scores and the dorsal pathway to BA44 for real noun morpho-syntax scores – a few additional findings were observed in only one of the two investigated samples. This highlights the need for replication of these findings in larger samples, which will be a significant challenge for future research given the difficulty of acquiring MRI data in young preschool children under 5 years of age. This would also help clarify which structures are involved in grammar before the age of 4. Further, in the present study we investigated grammar ability at the word-level, i.e. morpho-syntax. An exciting avenue for future research is the question of which age the association between grammar ability on the sentence-level and brain structural maturation of the dorsal pathway to BA44 arises, given the ongoing maturation of this pathway even during middle childhood (Brauer et al., 2013; Skeide et al., 2016). Moreover, here we tested children’s morpho-syntactic abilities with nouns. Given the different trajectories of nouns and verbs in language acquisition and the findings of dissociative neural correlates in adults for nouns versus verbs (Vigliocco et al., 2011; Waxman et al., 2013), future research should investigate whether the present findings also hold for the morpho-syntactic inflection of verbs.

### Conclusion

In sum, the present results highlight the role of the white matter language network for morpho-syntax acquisition during the critical preschool years between 3 and 5 years of age. The 4- and 5-year-olds, but not the 3-year-olds, showed a relationship between morpho-syntax and both the dorsal pathway to BA44 and the dorsal pathway to BA6. While in adults, the dorsal pathway to BA44 supports rule-based and the dorsal pathway to BA6 supports phonological processes, preschool-aged children seem to rely on both dorsal fiber pathways for morpho-syntax. Our findings suggest that young children may employ different strategies for acquiring the rule-based grammar system. The dorsal pathway to BA6 may support statistical learning mechanisms, whereas the maturation of the dorsal pathway to BA44 may support the acquisition of a robust rule-based grammar system.

## Acknowledgements

We thank Gesa Schaadt, Michael Skeide, Ting Qi and the entire LEGASCREEN consortium, and Hung Nguyen Trong and Christiane Attig for their support in data acquisition and sourcing. We thank Cornelius Eichner and Alfred Anwander for their methodological advice. We further thank Kerstin Flake, Andrea Gast-Sandmann and Heike Schmidt-Duderstedt for help with the figures, and Joshua Grant for helpful comments to a previous version of this manuscript. Further, we thank the Max Planck Förderstiftung for granting a writing stipend to C.C.K.

## Author Contributions

**Cheslie C. Klein**: Conceptualization, Data curation, Investigation, Methodology, Validation, Visualization, Formal analysis, Writing – Original Draft. **Philipp Berger**: Data curation, Methodology, Software, Supervision, Writing – Review & Editing. **Charlotte Grosse Wiesmann**: Conceptualization, Funding acquisition, Resources, Supervision, Investigation, Writing – Review & Editing. **Angela D. Friederici**: Conceptualization, Funding acquisition, Resources, Supervision, Writing – Review & Editing.

## Data availability statement

All employed materials and data discussed in the paper are saved in a local repository at the Max Planck Institute for Human Cognitive and Brain Sciences, and data in fully anonymized format will be made available to the reader upon request (according to the data protection policy in the ethics agreement).

