## Supplementary Material for "Grammar acquisition in preschool children is related to white matter maturation of the dorsal language network"

**Supplementary Table 1: Items of the morpho-syntactic word production task for each age group**

**Table 1:** Singular noun form of each item from the SETK 3-5 and its target plural form (Grimm, 2001), vowel change is indicated by capital letter.

|  | | **Singular form** | **Plural form** |
| --- | --- | --- | --- |
| (Practice item) | | Auto [engl. car] | Auto-s [engl. cars] |
| *3- to 5-y.o.s: Real nouns* | | |  |
|  | Fisch [engl. fish] | | Fisch-e [engl. fish] |
|  | Schiff [engl. ship] | | Schiff-e [engl. ships] |
|  | Gabel [engl. fork] | | Gabel-n [engl. forks] |
|  | Bild [engl. picture] | | Bild-er [engl. pictures] |
|  | Stuhl [engl. chair] | | StÜhl-e [engl. chairs] |
|  | Hand [engl. hand] | | HÄnd-e [engl. hands] |
|  | Buch [engl. book] | | BÜch-er [engl. books] |
|  | Glas [engl. glass] | | GlÄs-er [engl. glasses] |
|  | Vogel [engl. bird] | | VÖgel [engl. birds] |
|  | Apfel [engl. apple] | | Äpfel [engl. apples] |
| *4- to 5-y.o.s: Pseudo nouns* | | |  |
|  | Tulo | | Tulo-s |
|  | Biwo | | Biwo-s |
|  | Dolling | | Dolling-e |
|  | Ribane | | Ribane-n |
|  | Plarte | | Plarte-n |
|  | Tapsel | | Tapsel-n |
|  | Ropf | | Ropf-e/RÖpf-e |
|  | Kland | | KlÄnd-e |

**Supplementary Methods: Description of the plural noun system in German**

For plural formation, a word stem (e.g., dog) has to be combined with a plural morpheme (e.g., -s) to result in a plural word with an inherently more complex structure (e.g., dog-s) than the simple noun stem. In German, the plural of a noun can be externalized by several plural endings (i.e., –(e)n, –e, –s and –er) or no explicit marking on the noun, and additionally with a qualitative change of the stem vowel (e.g., Buch_neut_ – Büch-er [engl. book – book-s]; Werner, 1969). Thus, the German noun plural system consists of (multiple) rules depending primarily on noun phonology and gender (Köpcke, 1988). Most cases of plural nouns are covered by a few frequent rules (e.g., most female nouns take the plural marking –(e)n, as in Woche_fem_ – Woche-n [engl. week – week-s]; Köpcke, 1988). However, some irregular plural forms have to be lexicalized when no obligatory rule applies (e.g., the vowel change for masculine nouns with the plural –e, as in Ball_masc_ – Bäll-e [engl. ball – ball-s], but Tag_masc_ – Tag-e [engl. day – day-s]; Wegener, 1999). Frequent plural rules are typically acquired first during development and are also overapplied more often than less frequent rules (Kauschke, 2012).

### Supplementary Methods: Description of preregistered scores to assess children’s grammar ability on the sentence-level

We additionally preregistered two scores to assess children’s grammar ability on the sentence-level both in comprehension and production.

To receive the syntactic comprehension score, we used the subtest ‘Understanding sentences’ (orig. ‘Verstehen von Sätzen’, VS; Grimm, 2001) from the SETK 3-5 which consists of manipulation tasks. There, children are given instructions as, for example, to move objects in a certain order or to interact with another object. The items’ grammatical complexity of the presented sentence increases gradually, for instance by demanding causal relations between two actions and involving not only subject- but also object-initial sentences (VS: “Zeig mir: Der gelbe Ball, den der weiße Ball anstößt, fällt vom Tisch.” [engl. “Show me: The yellow ball bumped by the white ball falls off the table.”]; Grimm, 2001). While most items overlap in the tests for the 3- and 4- to 5-year-old children, some of the more complex constructions occur only in the older age groups as, for example, object-first subordinate clauses are not yet understood by young preschoolers (Grimm, 2001; Schipke et al., 2012). Additionally, the 3-year-olds participate in a sentence-picture-matching task in which they are asked to select one of four pictures matching the presented sentence. From this task, we then standardized the raw values within the respective age groups of each full behavioral sample, so within the 3- and 5-year-olds separately for Sample 1 (N = 194) and within Sample 2 of 3- and 4-year-olds (N = 58). These standardized values were then used as the syntactic comprehension score in further analyses.

To get an estimate for children’s syntactic production ability, we made use of the two sentence production subtests of the SETK 3-5, which were performed in the groups of 3- and 4- to 5-year-olds each. For the 3-year-old children, we used the production data from the SETK 3-5 subtest ‘Encoding semantic relations’ (orig. ‘Enkodierung semantischer Relationen’) which is a picture description task. On these pictures, people and animals perform an action with spatial relation to an object eliciting the use of prepositional phrases with varying degrees of difficulty depending on the required preposition (Grimm, 1975). For the 4- to 5-year-olds, we selected the subtest ‘Sentence memory’ (orig. ‘Satzgedächtnis’, SG), in which children are asked to reproduce sentences consisting of six to ten words with correct morpho-syntactic inflection and either plausible (SG: “Lena lacht, nachdem sie gekitzelt wurde.” [engl. “Lena laughs after being tickled.”]; Grimm, 2001) or implausible meaning (SG: “Ein frecher Fußball, der den alten Kasper heiratet, ist müde.” [engl. “A cheeky soccer ball marrying the old Punch is tired.”]; Grimm, 2001). The length of the sentences is determined, so that they cannot be solely retrieved from the child’s working memory but require reconstruction of the sentence structure using the child’s grammatical knowledge (Grimm, 2001). This effect is further enhanced by sentences with implausible meaning, as children cannot rely on their real-world knowledge (Grimm, 2001). To receive the syntactic production score, we recoded the production data to estimate the longest syntactically correct fragment in words per items as described in Klein et al. (2023). Then, we standardized these values within the age groups of each sample to account for differences in the respective tasks for 3- and 4- to 5-year-old children which were used in further analyses.

### Supplementary Methods: Description of preregistered score to assess children’s general language ability

To investigate the relation between children’s general language ability and brain structure, we preregistered an aggregated score from the SETK 3-5 as described in Klein et al. (2023). For this, we standardized the raw values of each subtest within the age groups of each sample to account for differences in the item structure of the test version for 3- versus 4- to 5-year-olds. Then, we combined these z-scores to the respective scale for language comprehension, production or memory as defined by the SETK 3-5 (Grimm, 2001) and summed these scale values to the general language score.

**Supplementary Table 2: Summary of the parameters for MRI acquisition in both samples**

**Table 2:** Scanning parameters used for acquisition of diffusion- and T1-weighted MRI data separated by samples, values for each parameter of the 3- & 5-year-old, and 3- & 4-year-old children.

| **Scanning parameter** | **Sample 1: 3- & 5-y.o.s** | **Sample 2: 3- & 4-y.o.s** |
| --- | --- | --- |
| **Head coil** | 12-channel | 32-channel |
| *Diffusion-weighted MRI* | | |
| **sequence** | optimized monopolar  Stejskal-Tanner EPI | multiplexed EPI |
| **TR** | 8,000 ms | 4,000 ms |
| **TE** | 83.0 ms | 75.4 ms |
| **b-value** | 1,000 s/mm2 | 1,000 s/mm2 |
| **directions** | 60 | 60 |
| **voxel size** | 1.86×1.86×1.9 mm | 1.9 mm isotropic |
| *T1-weighted MRI* | | |
| **sequence** | MP2RAGE | MP2RAGE |
| **TR** | 5,000 ms | 5,000 ms |
| **TE** | 2.82 ms | 3.24 ms |
| **TI_1_/TI_2_** | 700 ms/2,500 ms | 700 ms/2,500 ms |
| **α_1_/α_2_** | 4°/5° | 4°/5° |
| **voxel size** | 1.3 mm isotropic | 1.2×1.0×1.0 mm |

### Supplementary Figure 1: Masks used for segmentation to disentangle the two dorsal fiber pathways

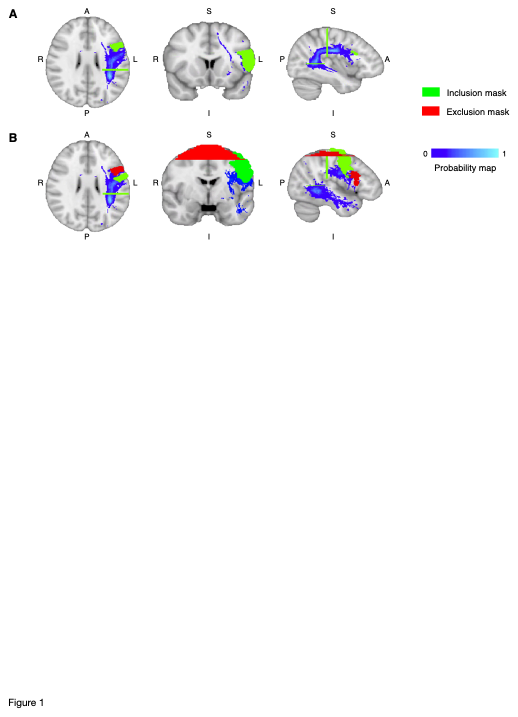

**Figure 1:** a) Masks used for segmentation of the dorsal fiber pathway targeting BA44 to include streamlines (green) based on prior anatomical assumptions. b) Masks used for segmentation of the dorsal fiber pathway targeting BA6 to include (green) and exclude (red) streamlines. Probability map of the arcuate fascicle (heat map) as provided by pyAFQ (Kruper et al., 2021) was used for refinement of both fiber pathways.

### Supplementary Figure 2: Examples for inclusion and exclusion criteria of the dorsal fiber pathways after manual quality check of segmentation results

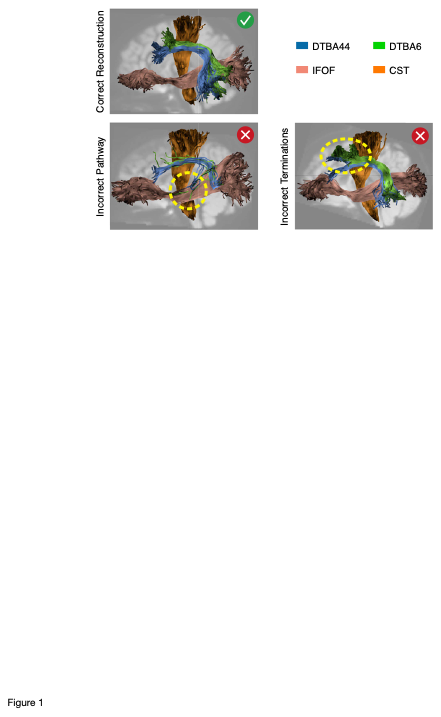

**Figure 2:** Inclusion criteria for segmentation of the two dorsal fiber pathways as the frontal part of both pathways terminate in distinct regions of BA44 and BA6, and two examples of excluded fiber pathways as streamlines were spurious or followed an anatomically implausible path, or frontal endpoints of pathways did not lead to distinct termination in the respective target regions of BA44 and BA6. Modified from Eichner, C., Berger, P., Klein, C. C., & Friederici, A. D. (2024). Lateralization of dorsal fiber tract targeting Broca’s area concurs with language skills during development. Progress in Neurobiology, 102602, https://doi.org/10.1016/j.pneurobio.2024.102602.

### Supplementary Figure 3: Overgeneralization errors of plural rules assigned to real and pseudo nouns in 3-, 4-, and 5-year-old children

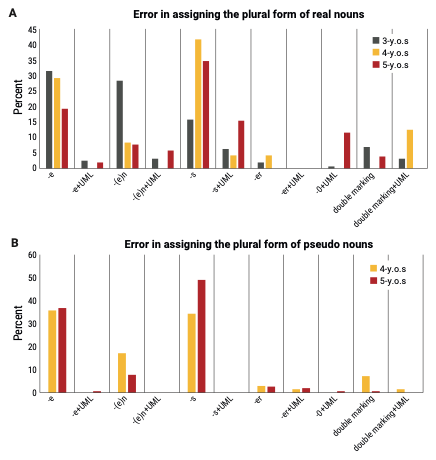

### Figure 3: Overapplication errors (in percent) of plural rules in preschool children aged 3, 4, and 5 years (N = 270). a) Percentage of errors in the assignment of plurals to real nouns (3-year-olds: gray; 4-year-olds: yellow; 5-year-olds: red). b) Percentage of errors in the assignment of plurals to pseudo nouns (4-year-olds: yellow; 5-year-olds: red).

### Supplementary Results: Preregistered analyses on the relation between preschoolers’ grammar ability on the sentence-level and language pathways

### We aimed to examine the relation between preschoolers’ grammar ability on the sentence-level and the maturation of language fiber pathways. For this, we preregistered two scores from a sentence comprehension and production task, and correlated them with children’s brain structural measures of the dorsal pathway to BA44, the dorsal pathway to BA6, the ventral pathway, and the corticospinal tract as a control. We controlled for sex, non-verbal IQ, handedness, eTIV, and, in Sample 1 of 3- and 5-year-olds, additionally for family history of dyslexia. Significance threshold was set at *p* = 0.05 for models with streamline count and for node-based models, multiple comparison correction was applied at *p* = 0.05 with a further Bonferroni-correction of N = 3.

### When testing for a relation between children’s syntactic comprehension scores and brain structure, we found no main effect in Sample 1 of 3- and 5-year-olds or Sample 2 of 3- and 4-year-olds in any language fiber pathway nor the control tract. Instead, we found a significant interaction with age group within both independent samples. In Sample 1, the 3- and 5-year-old children showed an age interaction in the dorsal pathway to BA44 (streamline count, β = -239.3, SE = 113.4, *t*(22) = -2.111, *p* = 0.04). No significant interaction with age group was found in the dorsal pathway to BA6, the ventral pathway, or the control tract in this sample. In Sample 2, the 3- and 4-year-olds also showed an interaction in the dorsal pathway to BA44 (anterior part: MD, node size = 19, node range = 1-19, β = -0.03, SE = 0.009, *t*(22) = -3.308, *p* = 0.003) and additionally in the ventral pathway (streamline count, β = 617.0, SE = 246.7, *t*(21) = 2.501, *p* = 0.02). No interaction was found in the dorsal pathway to BA6 or the corticospinal tract. We followed up on the present interactions and tested each age group separately. These analyses yielded no significant effect in any language fiber pathway or the control tract for either age group in both samples.

### We further tested for a correlation between children’s syntactic production scores and the maturation of language fiber pathways. We found no main effect when including both age groups of each sample and also no interaction with age group in neither sample.

### Supplementary Results: Preregistered analyses on the relation between preschooler’s general language ability and language pathways

In addition to our investigation on preschooler’s grammar ability, we further assessed the relation between their general language ability and the maturation of language fiber pathways. In a preregistered procedure, we related children’s overall performance in the general language test to brain structural measures in the three language pathways and the control tract in Sample 1 of 3- and 5-year-olds, and in Sample 2 of 3- and 4-year-olds, independently.

In neither sample, we found a significant main effect, including both age groups, in the language fiber pathways nor the control tract. We further found no significant interaction with age group in Sample 1. In Sample 2, we found a significant interaction with age group and children’s general language scores in the dorsal pathway to BA6 (central part: MD, node size = 28, node range = 40-67, β = -0.050, SE = 0.013, *t*(21) = -3.943, *p* < 0.001; RD, node size = 13, node range = 37-49, β = -0.089, SE = 0.021, *t*(21) = -4.282, *p* < 0.001) and the ventral pathway (streamline count, β = 936.9, SE = 364.7, *t*(20) = 2.569, *p* = 0.018), but not the dorsal pathway to BA44 or the corticospinal tract. When testing each age group separately to follow-up on this interaction, we found no significant effect in the 3- or 4-year-old children.

**Supplementary Results: Exploratory analyses on preschooler’s pseudo noun plurals and language pathways**

When testing for a relation between children’s pseudo noun morpho-syntax scores and brain structure, we found a significant relation with RD in the dorsal pathway to BA44 in the 4-year-olds in Sample 2 (anterior part: node size = 12, node range = 26-37, β = -0.027, SE = 0.006, *t*(11) = -4.664, *p* < 0.001). No effect was found in the dorsal pathway to BA6, the ventral pathway or the control tract. In Sample 1, the 5-year-old children showed no effect of their pseudo noun morpho-syntax scores in any language-related fiber pathway or the control tract.

### Supplementary Results: Exploratory analyses on preschooler’s morpho-syntactic ability and language pathways in the right hemisphere

In addition to our main analyses, we examined the relationship between children’s morpho-syntax scores and brain measures of the three homologues language pathways (i.e., the dorsal pathway to BA44 and BA6, and IFOF) and control tract (i.e., the corticospinal tract) in the right hemisphere. This was done to test whether preschoolers exhibit hemisphere-specific differences in the association between brain structures and morpho-syntactic ability, given the left-lateralization of language in the adult brain (Toga & Thompson, 2003).

Consistent with the findings in the left hemisphere, no significant main effect was observed between children’s morpho-syntax scores and brain measures in the right fiber pathways in either of the two samples. Instead, there was a significant interaction with age group in both samples.

In Sample 1 of 3- and 5-year-old children, the age interaction was found in the right ventral pathway (streamline count, β = 568.2, SE = 158.4, *t*(80) = 3.588, *p* < 0.001; central part: FA, node size = 14, node range = 31-44, β = 0.030, SE = 0.007, *t*(80) = 4.236, *p* < 0.001). In contrast to the findings in the left hemisphere, no age interaction was found in the homologous right dorsal pathway to BA6. Further, we found no interaction in the right dorsal pathway to BA44 similar to the prior analyses in the left hemisphere. Analyses by age group revealed that the interaction was driven by the 5-year-old children who showed a significant relation between morpho-syntax scores and the right ventral pathway (streamline count, β = 408.9, SE = 132.5, *t*(48) = 3.085, *p* = 0.002), but not with any of the two dorsal pathways. 3-year-olds did not show a significant effect in any of the right language pathways, paralleling the findings in the left hemisphere.

In Sample 2 of 3- and 4-year-old children, we again found no main effect in the right-hemispheric language pathways, but a significant interaction with age group. This age interaction was observed in the right dorsal pathway to BA6 (anterior part: RD, node size = 16, node range = 8-23, β = -0.058, SE = 0.015, *t*(17) = -3.843, *p* = 0.001) similar to the results in the left hemisphere. However, in contrast to the prior results, we did not find an interaction in the right dorsal pathway to BA44, but instead in the right ventral pathway (posterior part: MD, node size = 26, node range = 63-88, β = -0.045, SE = 0.010, *t*(22) = -4.376, *p* < 0.001; RD, node size = 13, node range = 70-82, β = -0.058, SE = 0.014, *t*(22) = -4.248, *p* < 0.001) replicating the findings of Sample 1. Further analyses revealed that the interaction was driven by the 4-year-old children, while the 3-year-olds showed no significant relation. These findings are consistent with the results observed in the left hemisphere of the same children and replicate the results from Sample 1. Conversely, the 4-year-old children showed a significant relation between their morpho-syntax scores and the right dorsal pathway to BA6 (anterior part: MD, node size = 36, node range = 7-36, β = -0.029, SE = 0.003, *t*(8) = -9.330, *p* < 0.001; RD, node size = 24, node range = 4-27, β = -0.029, SE = 0.004, *t*(8) = -6.672, *p* < 0.001). However, no significant association was found with the right ventral pathway, contrary to the observed interaction. Additionally, in contrast to the left hemisphere findings, we found no relation with the right dorsal pathway to BA44 in the 4-year-olds.

No significant main effect or interaction was found for the control tract (i.e., the right corticospinal tract) in either sample.

### Supplementary Results: Exploratory analyses on preschooler’s real noun plurals and language pathways in the right hemisphere

For children’s morpho-syntax scores tested only with real nouns, again no main effect was observed in the right-hemispheric language pathways in either sample. However, a significant interaction with age group was found, in line with the results in the left hemisphere.

In Sample 1 of 3- and 5-year-olds, the interaction was found in the dorsal pathway to BA6 (central part: MD, node size = 20, node range = 30-46, β = -0.032, SE = 0.009, *t*(71) = -3.460, *p* < 0.001), in the dorsal pathway to BA44 (central part: MD, node size = 15, node range = 43-57, β = -0.028, SE = 0.008, *t*(74) = -3.591, *p* < 0.001), and the ventral pathway (streamline count, β = 630.9, SE = 194.3, *t*(79) = 3.248, *p* = 0.002; central part: FA, node size = 15, node range = 32-46, β = 0.034, SE = 0.008, *t*(79) = 4.186, *p* < 0.001; RD, node size = 14, node range = 32-45, β = -0.042, SE = 0.012, *t*(79) = -3.642, *p* < 0.001). As before, follow-up analyses revealed that these interactions were driven by the older age group. Specifically, the 5-year-olds showed a significant effect in the right ventral pathway (streamline count, β = 440.3, SE = 185.8, *t*(48) = 2.370, *p* = 0.011; anterior to central part: FA, node size = 11, node range = 35-45, β = 0.022, SE = 0.007, *t*(48) = 2.977, *p* = 0.002). No effect, however, was found for the dorsal pathway to BA6 or BA44 in the older age group, which contrasts with the left-hemispheric findings of a relation with both dorsal pathways.

In Sample 2 of the 3- and 4-year-old children, the interaction was also located in the dorsal pathway to BA6 (anterior to central part: RD, node size = 12, node range = 33-44, β = -0.069, SE = 0.017, *t*(17) = -4.067, *p* < 0.001). This replicates the left-hemisphere findings in the same children and aligns with the results of Sample 1. No significant interaction, however, was found in the dorsal pathway to BA44 or the ventral pathway in this sample. Following-up on this interaction, 4-year-old children did not show a significant effect in the dorsal pathway to BA6, contrary to the observed interaction. Further, no effect was found in the right dorsal pathway to BA44 or the right ventral pathway in the 4-year-olds.

As in the prior analyses, we found no significant effect for 3-year-olds’ real noun morpho-syntax scores in either sample. Moreover, no main effect or interaction was found for the right-hemispheric control tract in either sample.

### Supplementary Results: Exploratory analyses on preschooler’s pseudo noun plurals and language pathways in the right hemisphere

When testing for a relation between children’s morpho-syntax scores tested only with pseudo nouns and brain structure in the right hemisphere, we found an effect in the ventral pathway (streamline count, β = 240.2, SE = 115.0, *t*(47) = 2.089, *p* = 0.021) in the older age group of Sample 1, i.e. the 5-year-olds. No effect was found in the right dorsal pathway to BA44 or BA6, or the control tract in this age group.

In Sample 2, the 4-year-old children showed an effect for their pseudo noun morpho-syntax scores in the right dorsal pathway to BA6 (anterior part: MD, node size = 31, node range = 9-39, β = -0.033, SE = 0.004, *t*(8) = -7.384, *p* < 0.001; RD, node size = 22, node range = 5-26, β = -0.034, SE = 0.005, *t*(8) = -6.843, *p* < 0.001). In contrast to the findings in the left hemisphere, the 4-year-olds did not show a significant relation with their right dorsal pathway to BA44. Further, no effect was found for the right ventral pathway or the control tract.
